## Supplementary materials for "ATP-release pannexin channels are gated by lysophospholipids"

Erik Henze[1†], Russell N. Burkhardt[2,3], Bennett W. Fox[2,3], Tyler J. Schwertfeger[2,3], Eric Gelsleichter[3], Kevin Michalski[1], Lydia Kramer[1‡], Margret Lenfest[4], Jordyn M. Boesch[4], Hening Lin[3,5,6§], Frank C. Schroeder[2,3], and Toshimitsu Kawate[1*]
[1]Department of Molecular Medicine, Cornell University, Ithaca, NY 14853, USA
[2]Boyce Thompson Institute, Cornell University, Ithaca, NY, USA.
[3]Department of Chemistry and Chemical Biology, Cornell University, Ithaca, NY, USA.
[4]Department of Clinical Sciences, College of Veterinary Medicine, Cornell University, Ithaca, NY, USA.
[5]Department of Molecular Biology and Genetics, Cornell University, Ithaca, NY, USA.
[6]Howard Hughes Medical Institute, USA

*Corresponding author:

Toshimitsu Kawate
Department of Molecular Medicine
Cornell University
C4-151 VMC, 930 Campus Rd, Ithaca, NY 14853


The supplementary information includes: Figures S1 to S5, Table S1

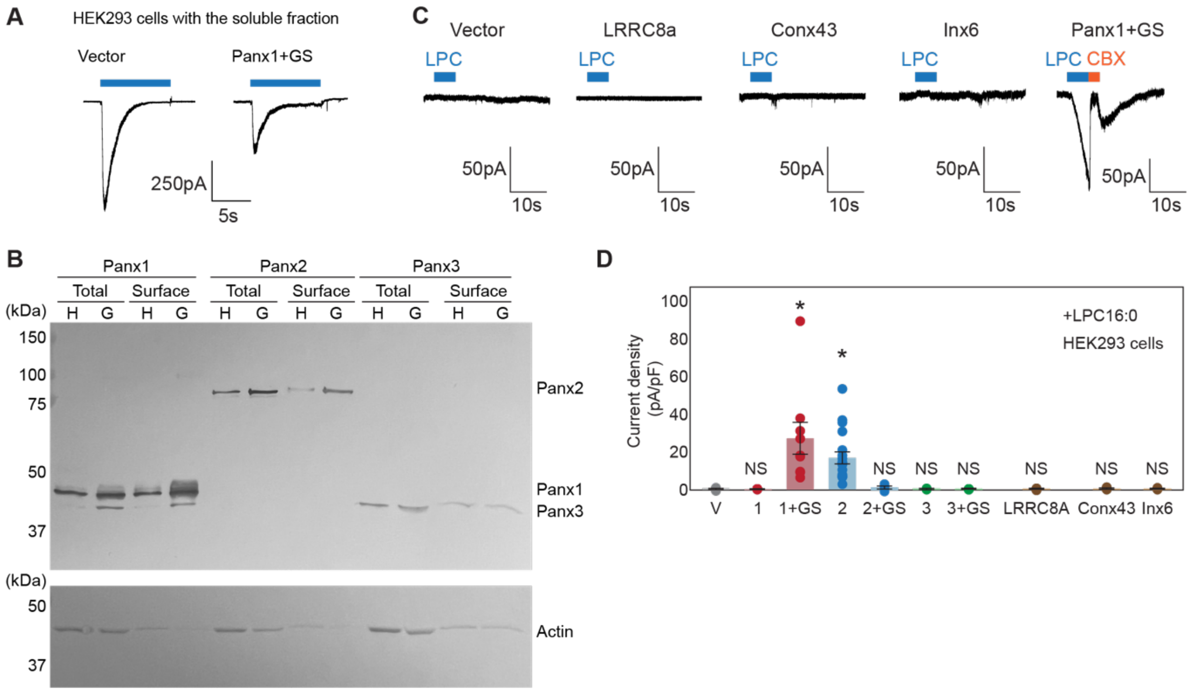


**Fig. S1: Characteristics of the pannexins and other large pore forming channels used in this study.** (A) Representative whole-cell patch-clamp traces with the soluble fraction from mouse liver extract. Vector (left) or Panx1+GS (right) were expressed in HEK293 cells and voltage clamp recordings were performed at -60 mV. The blue bars indicate application of the soluble fractions. (B) Cell surface biotinylation pulldown to determine surface protein levels of heterologously-expressed pannexins in HEK293 (H) and HEK293S GnTI$^-$ (G) cells. Representative western blots for the total and surface biotinylated samples are shown. Top: anti-FLAG; bottom; anti-actin. (C) and (D) Whole-cell patch-clamp recordings of various constructs used in this study. Representative whole-cell patch-clamp traces (C) and quantification of the peak currents obtained with LPC-16:0 (7 μM). Pannexins and other members of the large-pore channel family are expressed in HEK293 cells and voltage clamp recordings were performed at -60 mV. Blue bars indicate application of LPC-16:0 and red bars indicate application of CBX (50 μM). N=4-19. P values are obtained using one-way ANOVA followed by Dunnett's t-test. Asterisks indicate $P<0.01$ and error bars represent s.e.m.

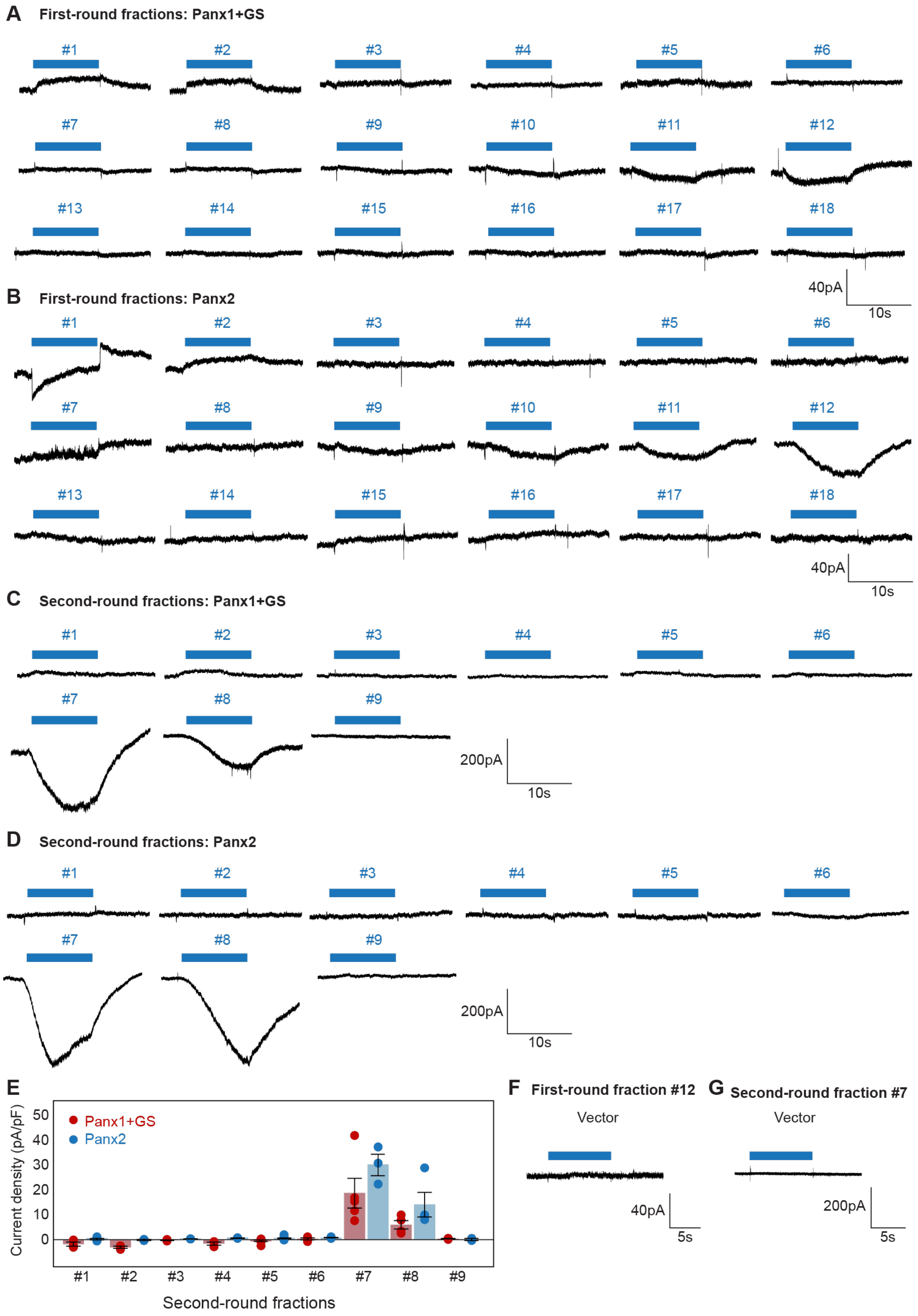
A First-round fractions: Panx1+GS
#1 #2 #3 #4 #5 #6
#7 #8 #9 #10 #11 #12
#13 #14 #15 #16 #17 #18
40pA
10s
B First-round fractions: Panx2
#1 #2 #3 #4 #5 #6
#7 #8 #9 #10 #11 #12
#13 #14 #15 #16 #17 #18
40pA
10s
C Second-round fractions: Panx1+GS
#1 #2 #3 #4 #5 #6
#7 #8 #9
200pA
10s
D Second-round fractions: Panx2
#1 #2 #3 #4 #5 #6
#7 #8 #9
200pA
10s
E
Current density (pA/pF)
50 40 30 20 10 0
Panx1+GS
Panx2
#1 #2 #3 #4 #5 #6 #7 #8 #9
Second-round fractions
F First-round fraction #12
Vector
40pA
5s
G Second-round fraction #7
Vector
200pA
5s

**Fig. S2: Representative whole-cell patch-clamp traces for mouse liver fractions.** (A-D) Pannexins were expressed in HEK293 cells and voltage clam recordings were performed at -60 mV. Both the first round ((A) and (B)) and the second round ((C) and (D)) fractions are shown. Blue bars indicate application of each fraction. (E) Current densities from second-round fractions. N=3-5. Error bars represent s.e.m. (F) and (G) Whole-cell currents obtained with vector-transfected control cells. Application of fraction #12 from the first round and fraction #7 from the second round are shown.

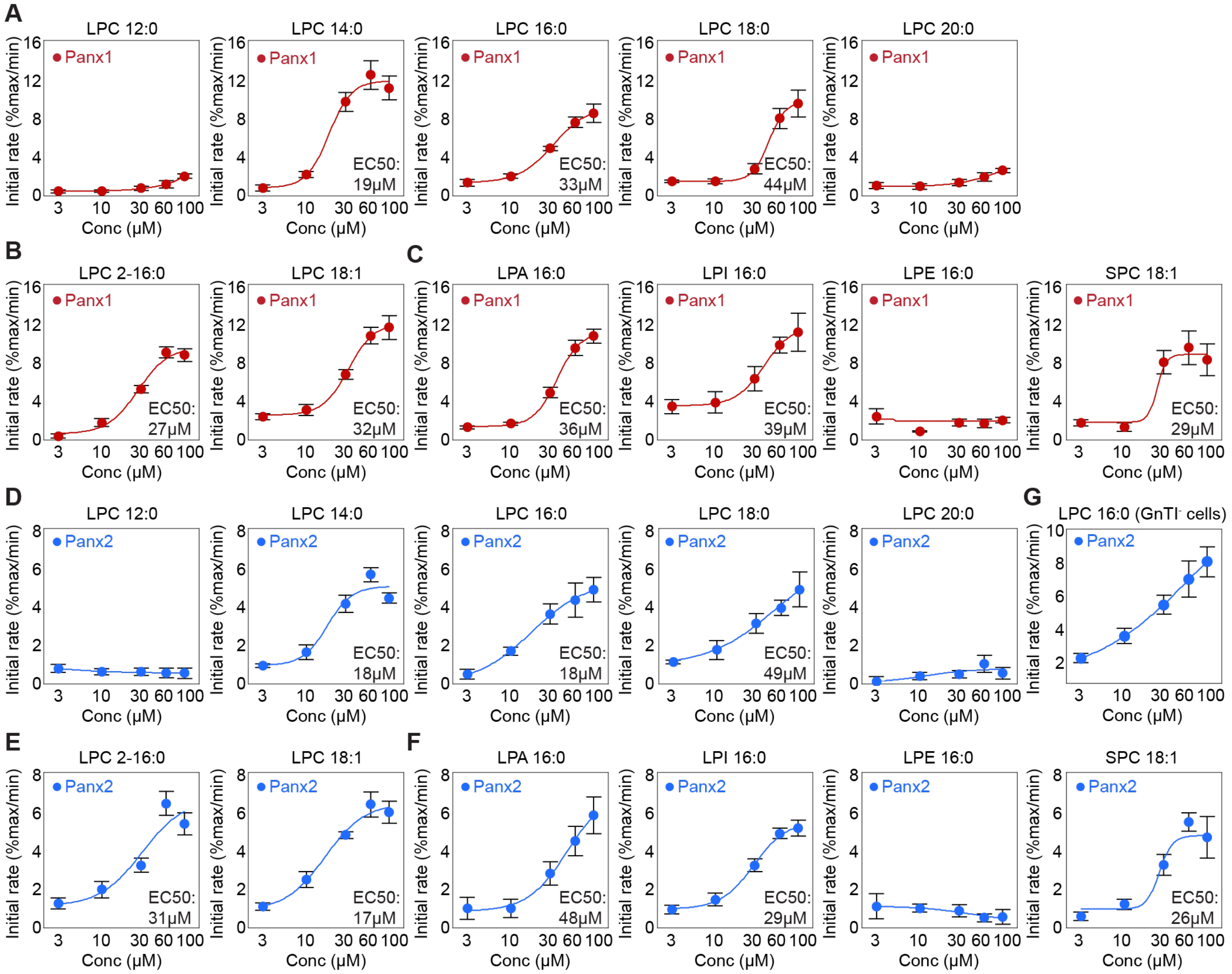


**Fig. S3: Lysophospholipid dose-response curves obtained from the mVenus quench assay.** (A-C) Panx1 dose-response curves for LPC with different chain length (A), LPC with different tail branching (B), and lysophospholipids with different headgroups (C). Wildtype Panx1 was expressed in GnTI$^-$ cells. (D-G) Wildtype Panx2 dose-response curves obtained from HEK293 cells for LPC with different chain length (D), LPC with different tail branching (E), lysophospholipids with different headgroups (F), and with LPC16:0 in GnTI$^-$ cells (G). Fluorescence of the vector control was subtracted from the total activity. Mean of at least four independent experiments are shown. Dose responses were fitted with the Hill equation and the EC50 values are indicated. Error bars represent s.e.m.

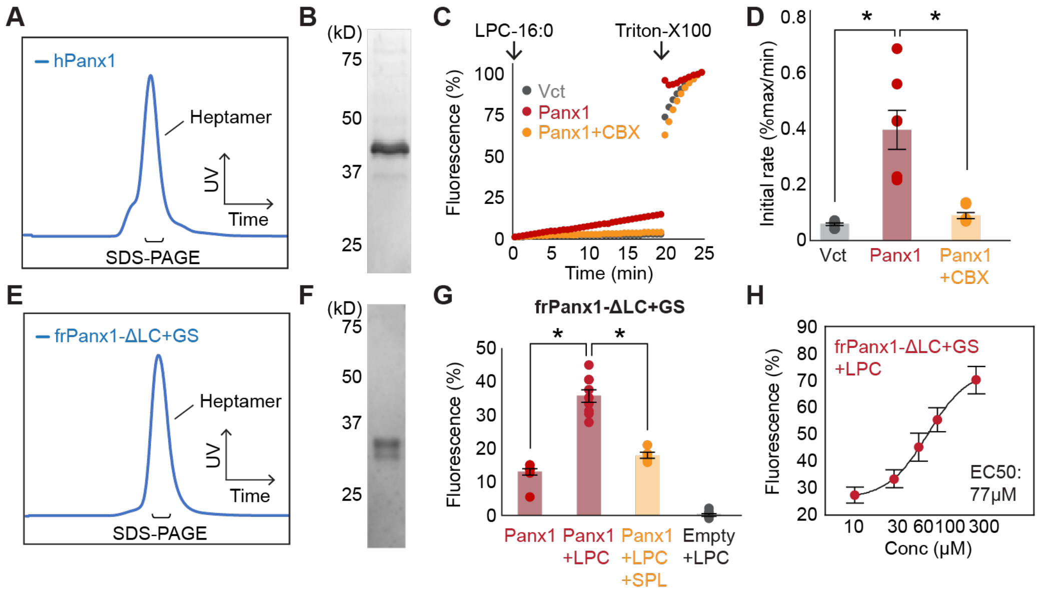


**Fig. S4: Functional reconstitution of purified Panx1.** (A-D) Full-length human Panx1 used for functional reconstitution. A representative SEC profile (A) and an SDS-PAGE gel image (B) of the samples purified from HEK293S GnTI$^-$ cells. Time course (C) and quantification (D) of cellular YO-PRO-1 uptake are shown. LPC-16:0 (10 μM) and CBX (50 μM) are used. N=8. P values are obtained using one-way ANOVA followed by Dunnett's t-test. Asterisks indicate P<0.01 and error bars represent s.e.m. (E-H) The frog Panx1 construct (frPanx1-ΔLC+GS) used for the previous cryo-EM studies *(32)*. A representative SEC profile (E) and an SDS-PAGE gel image (F) of frPanx1-ΔLC+GS purified from High Five insect cells. Relative YO-PRO-1 fluorescence from frPanx1-ΔLC+GS reconstituted liposomes triggered by LPC16:0 (100 μM) (G). Spironolactone (SPL; 50 μM) was used to confirm Panx1-specific activity. P values are obtained using unpaired t-test. Asterisks indicate P< 0.01. N=6-15. Dose-response profile of Panx1 treated with LPC-16:0 (H). N=9-11. Dose responses were fitted with the Hill equation and the EC50 values are indicated. Error bars represent s.e.m.

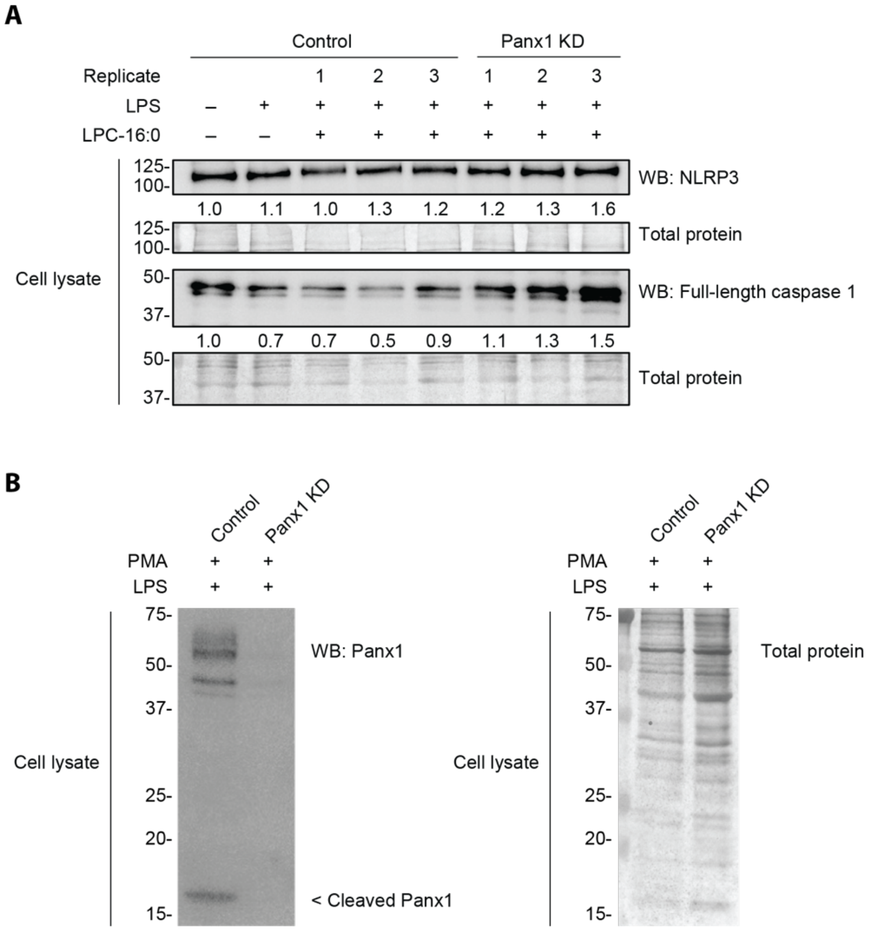


**Fig. S5: Expression of Panx1 and other inflammasome components in THP-1 cells treated with LPC16:0.** (A) Protein expression levels of NLRP3, full-length caspase 1, and (B) Panx1 in PMA-differentiated/LPS-primed THP-1 cells used in LPC16:0-induced IL-1β release experiments. Relative NLRP3 and caspase 1 levels (A) in cell lysates were quantified with densitometry by normalizing to total protein loading.

### Table S1

**Enriched compounds in fractions 7 and 8**

| RT (min) | [M]+ | Obs. *m/z* | Calc. *m/z* | Error (ppm) | [M]- | Obs. *m/z* | Calc. *m/z* | Error (ppm) | Formula | ID |
|---|---|---|---|---|---|---|---|---|---|---|
| 15.05 | ND | - | - | - | M-H | 229.1808 | 229.1809 | 0.46 | C13H26O3 | Unknown. Likely hydroxylated fatty acid |
| 15.1 | M+H | 454.2917 | 454.2928 | 2.37 | M-H | 452.2782 | 452.2783 | 0.24 | C21H44NO7P | LPE(0/16:0)* |
| 15.12 | ND | - | - | - | M-H | 243.1964 | 243.1966 | 0.54 | C14H28O3 | Unknown. Likely hydroxylated fatty acid |
| 15.16 | M+H | 335.3047 | 335.3057 | 3.09 | ND | - | - | - | C21H38N2O | Unknown |
| 15.25 | ND | - | - | - | M-H | 241.1809 | 241.1809 | 0.06 | C14H26O3 | Unknown. Likely hydroxylated fatty acid |
| 15.32 | ND | - | - | - | M-H | 243.1964 | 243.1966 | 0.51 | C14H28O3 | Unknown. Likely hydroxylated fatty acid |
| 15.38 | M+H | 454.2918 | 454.2928 | 2.27 | M-H | 452.2781 | 452.2783 | 0.30 | C21H44NO7P | LPE(16:0/0) |
| 15.39 | M+H | 496.3388 | 496.3398 | 1.85 | M+formate | 540.3296 | 540.3307 | 2.03 | C24H51NO7P | LPC(0/16:0) |
| 15.56 | M+H | 546.3548 | 546.3554 | 1.15 | M+formate | 590.3465 | 590.3463 | -0.21 | C28H53NO7P | LPC(20:3/0)* |
| 15.68 | M+H | 496.3389 | 496.3398 | 1.80 | M+formate | 540.3297 | 540.3307 | 1.89 | C24H51NO7P | LPC(16:0/0) |
| 15.76 | M+H | 522.3548 | 522.3554 | 1.25 | M+formate | 566.3469 | 566.3463 | -0.96 | C26H53NO7P | LPC(0/18:1)* |
| 16.01 | M+H | 522.3546 | 522.3554 | 1.65 | M+formate | 566.3468 | 566.3463 | -0.86 | C26H53NO7P | LPC(18:1/0) |
| 21.86 | M+H (?) | 536.1646 | - | - | ND | - | - | - | ? | Unknown** |

ND; not detected
* Predicted metabolite based on molecular formula and MS/MS fragmentation.
**Likely polysiloxane or similar contaminant. Detected most strongly in fraction 8, but additionally in fractions 4 and 5.
